# Preferential Invasion of Differentiated Bladder Carcinoma Cells by Flagellated Group B2 *Escherichia Coli*

**DOI:** 10.64898/2026.04.16.718932

**Authors:** Jacob Hogins, Julia Nguyen, Waris Khuwaja, Sydney Hall, Virginia Fogg, Xintong Dong, Philippe Zimmern, Larry Reitzer

## Abstract

Phylogenetic group B2 *Escherichia coli* is associated with urinary tract infections and other pathologies, but the basis for this phylogenetic skew is not understood. One aspect of urinary tract infections is binding to and entering uroepithelial cells. To test whether a phylogenetic skew exists for cell invasion, we examined invasion of 10 *E. coli* strains from three phylogenetic groups into CRL2169 and HTB-9 cells, which are derived from grade 1 and grade 2 bladder carcinomas, respectively. The top four strains that invaded CRL2169 were from group B2: three of these strains had more flagella gene transcripts than the other seven strains. The seven strains that invaded HTB-9 were from different phylogenetic groups. For the model uropathogenic group B2 strain UTI89, which expresses pili over flagella, loss of flagella or pili impacted invasion into CRL2169 to similar extents, but loss of pili had a greater effect on invasion into HTB-9 and a murine infection model than loss of flagella. A hyperflagellated variant of a group A strain did not invade either cell line better than the parental strain. Reported transcript differences, which were confirmed experimentally, showed that CRL2169 was more differentiated. The endocytosis stimulator tanshinone enhanced invasion into HTB-9, but not into CRL2169, which suggests differences in endocytic pathways and is consistent with differences in differentiation states. If the initial or recurring event in urinary tract infection is invasion into differentiated urothelial cells, as opposed to tight junctions, then the role of flagella may have been underestimated.

## Introduction

Urinary tract infections (UTIs) are the most common community-acquired bacterial infection in the United States (1, 2). UTIs occur more often in women and their frequency increases with age (3, 4). Without treatment, these infections can last years, leading to heavy financial, physical, and psychological burdens and can result in possible loss of the bladder (5). Antibiotics are the frontline treatment method for UTIs, but invasive intracellular bacteria are not susceptible to antibiotics. A prolonged infection and the associated treatment can lead to antibiotic resistance. The World Health Organization recognizes the bladder during a UTI as a major source of developing multidrug resistance (6).

Studies with model uropathogenic organisms have shown that infections require ascension to the bladder from the urethra; rapid growth in the nutrient-limited bladder lumen; attachment to and entry into uroepithelial cells; rapid intracellular replication and release; and resistance to innate immunity factors (1). Uropathogenic *Escherichia coli* (UPEC) is most often associated with UTIs (7). *E. coli* is a heterogenous species with multiple phylogenetic groups (8–11): group B2 strains account for 65-75% of *E. coli*-based UTIs (7, 12, 13). The basis for the group B2 phylogenetic skew is not understood. A few B2-specific genes or alleles have been implicated in virulence (14, 15), but group A and D strains are also pathogenic which implies that the B2-specific genes are not absolutely necessary. In addition to differences in genes, ∼50% of group B2 core genes are differentially expressed compared to group A and D strains when grown in a nutrient-rich medium (16). The core gene transcriptomes of pathogenic and non-pathogenic strains are not obviously different within the phylogenetic groups (16).

One component of a UTI is binding to and invasion into uroepithelial cells which are complex processes that appear to involve multiple pathways (17). The model UPEC strain UTI89 relies on type 1 pili to invade both bladder and vaginal epithelial cells (18, 19). Adhesion requires the pilus protein FimH which binds the mannosylated proteins β1α3 integrin and the uroplakins (17, 20, 21). Binding activates an endocytosis pathway known as the zipper pathway which requires dynamin, caveolin-1 coated vesicles, and Src signaling in the human cell (21, 22). Several FimH variants exist, and specific variants are associated with virulence (15, 23, 24). Deletion of the *fim* operon, which codes for pili components, does not abolish invasiveness, which implies an alternative entry mechanism (19). One flagella variant has been shown to contribute to intracellular invasion and immune evasion which implicates flagella in binding to the epithelium (25–27). Flagella bind to the human toll-like receptor 5 (TLR5) and trigger cell invasion that involves dynamin, clathrin-coated vesicles, and the NF-κB pathway (28). In addition, lipopolysaccharide has been indirectly implicated in epithelial binding since specific toll-like receptor 4 (TLR4) variants have been associated with UTIs in Asian populations (29), and lipopolysaccharide binds TLR4 (30).

Much of our knowledge of these processes is dependent on model UPEC strains, such as UTI89, which is not representative of other B2 strains (16), and a single human male bladder cell line (HTB-9), which is derived from a dedifferentiated class 2 bladder carcinoma. We examined 10 sequenced bacterial strains from three phylogenetic groups for invasion into HTB-9 and the CRL2169 cell line, the latter of which is derived from a relatively differentiated class 1 bladder carcinoma of a female patient. The differentiation state is important because of effects on the endocytic pathways: endocytosis is minimal in healthy uroepithelial umbrella cells, and, conversely, endocytosis increases during oncogenic dedifferentiation (31, 32). We show a phylogenetic skew towards B2 strains for invasion into CRL2169, but not into HTB-9, and that invasion into CRL2169 involves flagella. We conclude that the contribution of pili or flagella for invasion depends on properties of the epithelial cells, which has implications for the mechanism that initiates a UTI.

## Results

### A group B2 phylogenetic skew for invasion into CRL2169 but not into HTB-9

We analyzed invasion, which requires adhesion, endocytosis, and intracellular maintenance, for 10 strains of *E. coli* that represented three phylogenetic groups into HTB-9 and CRL2169 which are derived from grade 2 and grade 1 carcinomas, respectively. Table 1 lists the bacterial strains, their phylogenetic group and sequence type, whether they were isolated from an individual with a UTI, and summarizes the invasiveness of the bacterial strains.

**Table 1.** Summary of strains and their properties.

| Strain | Phylogenetic Group | Sequence Type | UPEC<sup>a</sup> | Pili (P) or Flagella (F) Dominant<sup>b</sup> | HTB-9 invasion<sup>c</sup> | CRL2169 invasion<sup>c</sup> | Source |
| --- | --- | --- | --- | --- | --- | --- | --- |
| ECOR11 | A | 10 | Yes | Neither | + | — | (55) |
| ECOR14 | A | 10 | Yes | F (P) | + | — | (55) |
| W3110 | A | 10 | No | P | — | — | Lab strain |
| W3110-J1 | A | 10 | No | F | — | — | (34) |
| ECOR44 | D | 66 | No | P | + | — | (55) |
| ECOR48 | D | 70 | Yes | P | + | — | (55) |
| CFT073 | B2 | 73 | Yes | F | + | + | (58) |
| ECOR51 | B2 | 73 | No | F | + | + | (55) |
| ECOR53 | B2 | 12 | No | P | — | — | (55) |
| LRPF007 | B2 | 95 | Yes | F | — | + | (56) |
| UTI89 | B2 | 95 | Yes | P | + | + | (57) |
| UTI89 $\Delta fimA$ | B2 | 95 | NA | F | — | + | This work |
| UTI89 $\Delta fliC$ | B2 | 95 | NA | P | — | -/+ | This work |
| UTI89 $\Delta fimA \Delta fliC$ | B2 | 95 | NA | NA | — | — | This work |

a. UPEC (uropathogenic *E. coli*) is defined as isolated from a urinary tract infection patient. Transcriptomic results suggest that pathogenic and non-pathogenic are indistinguishable (16). Since *E. coli* is an opportunistic pathogen, even the non-pathogenic strains may be virulent.
b. The designation is based on our transcriptomic analysis of these strains (16)
c. Invasiveness is defined as $\geq 50$ bacteria isolated from the invasion cultures.

For invasion into the well-studied HTB-9, seven bacterial strains from three phylogenetic groups had > 50 cells recovered (Fig 1A). The five B2 strains collectively invaded slightly better (p = 0.039), but the ranges from strains of group B2 and groups A and D had substantial overlap (Fig 1B). For invasion into CRL2169, the four most invasive strains (>50 cells recovered) were the group B2 strains CFT073, UTI89, LRPF007, and ECOR51 (Fig 1C and D), and each invaded better than any group A or D strain. Group B2 strains CFT073, UTI89, and ECOR51 invaded both cell lines equally well whereas group A and D strains invaded HTB-9 cells better than CRL2169 cells (Fig 1E and F). LRPF007 (a recently isolated group B2 strain from a UTI patient) was the only strain that invaded CRL2169 better than HTB-9.

**Figure 1.**
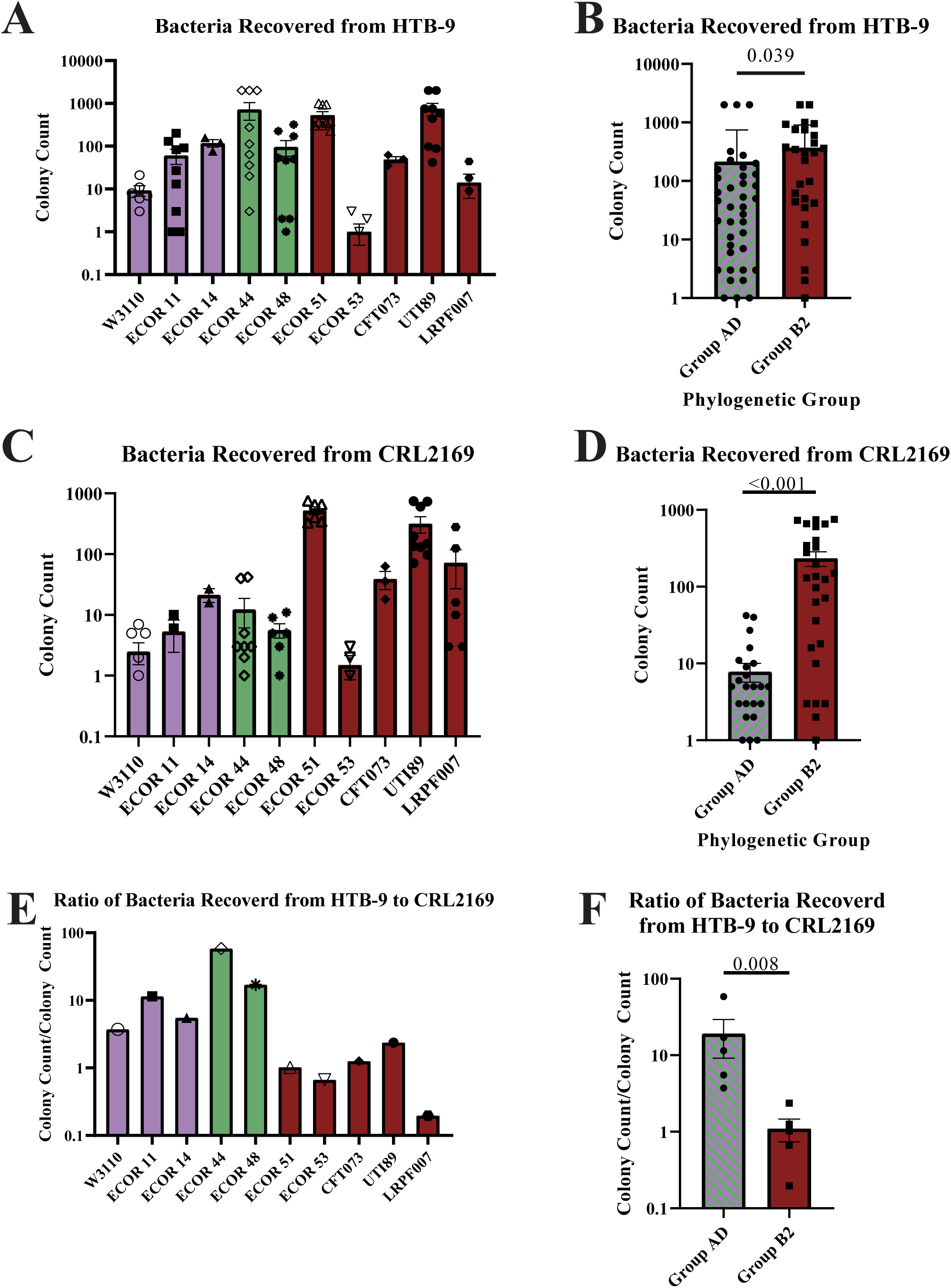
Invasion of *E. coli* strains from three phylogenetic groups in HTB-9 and CRL2169. Group AD refers to strains from either phylogenetic group A or D. IBC (intracellular bacterial community) refers to intracellular bacteria recovered. (A) Invasion into HTB-9: colonies recovered. (B) Invasion into HTB-9: cumulative comparison of group B2 vs non-B2 strains. This panel replots all the technical replicates for each strain from panel A. (C) Invasion into CRL2169: colonies recovered. (D) Invasion into CRL2169: cumulative comparison of group B2 vs non-B2 strains. This panel replots all the technical replicates for each strain from panel C. (E) Ratio of invasion into HTB-9 to CRL2169: individual strains. (E) Ratio of invasion into HTB-9 to CRL2169: cumulative comparison.

We previously analyzed the transcriptome of 35 strains that had been grown in a glucose-containing nutrient-rich medium which generally describes the invasion medium (16). An updated graphic representation of transcriptomic groups for 42 strains is shown in Fig. A1. The transcriptomes cluster into B2 and non-B2 transcriptome types, although some B2 strains have the non-B2-type transcriptome and vice versa. Of the four strains that invaded CRL2169, which are all group B2 strains, ECOR51 and LRPF007 had the B2-type transcriptome and CFT073 and UTI89 had the non-B2-type transcriptome. In summary, a group B2 phylogenetic skew was observed for invasion into CRL2169 but not into HTB-9.

### Flagella are important for invasion into CRL2169

Our transcriptomic analysis showed substantial strain variation for pili and flagella expression (Fig 2A and B). From the transcript ratio of *fimA* to *fliC*, which code for the major structural subunits of pili and flagella, respectively, we classify the strains as either pili- or flagella-dominant (Fig 2C). Of the four strains that invaded CRL2169 (>50 cells recovered), ECOR51, CFT073, and LRPF007 are flagella-dominant, while UTI89 is pili-dominant. Of the seven strains that invaded HTB-9, three are flagella-dominant (ECOR14, ECOR51, and CFT073); three are pili-dominant (ECOR44, ECOR48, and UTI89); and ECOR11 poorly expressed pili and flagella.

**Figure 2.**
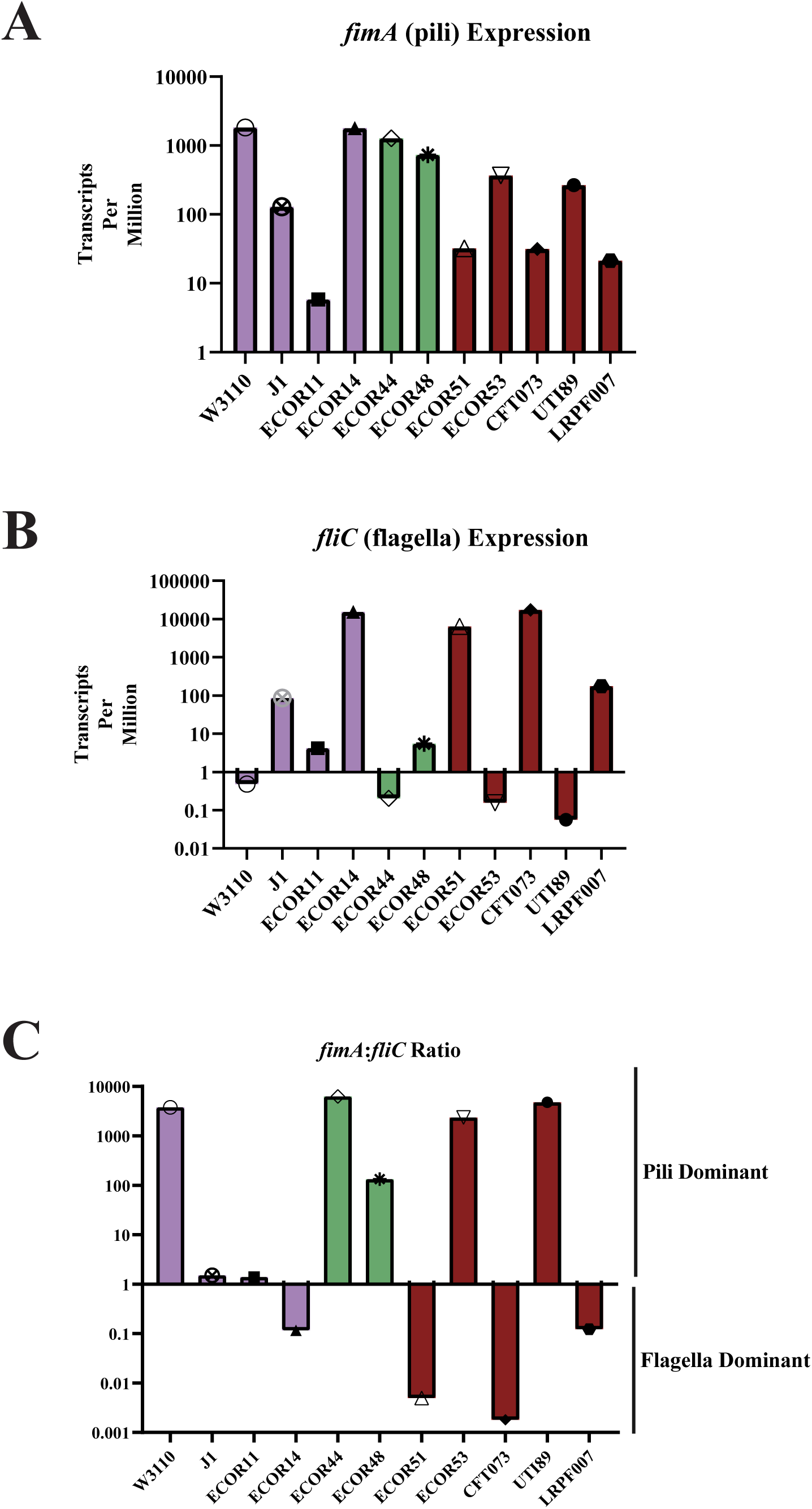
Pili (*fimA*) and flagella (*fliC*) structural subunits expression in the *E. coli* strains. TPM data was taken from the NCBI depository and had been generated by the authors. (A) *fimA* transcripts; (B) *fliC* transcripts; (c) ratio of *fimC* to *fliC* transcripts.

We examined how loss of flagella and pili in pili-dominant UTI89 affected invasion into CRL2169 and HTB-9 and in a mouse infection model. For invasion into CRL2169 cells, loss of *fliC* reduced invasion 75% (p = 0.074), and loss of *fimA* reduced invasion by 25% (p = 0.865) (Fig 2A). Loss of flagella appears to be modestly more impactful. Loss of both appendages effectively eliminated invasion (Fig 3A). For invasion into HTB-9, loss of *fliC* reduced invasion 80% (p = 0.031), and loss of *fimA* decreased invasion by 90% (p = 0.05) (Fig 3B). The ranges for the Δ*fimA* and Δ*fliC* strains did not overlap with the range for the parental strain which suggests that loss of either appendage is meaningful. The double mutant did not invade (Fig 3B). In a well-established transurethral mouse infection model (33), loss of *fimA* reduced recoverable bacteria by 99% (p = <0.0001), loss of *fliC* reduced the bacterial count by 50% (p = 0.093), and loss of both genes rendered the bacteria unable to infect the mouse bladder (Fig 3C). In summary, loss of flagella was more important than loss of pili for invasion into CRL2169, but not for invasion into HTB-9 or a mouse infection, and only loss of both appendages led to complete non-invasiveness.

**Figure 3.**
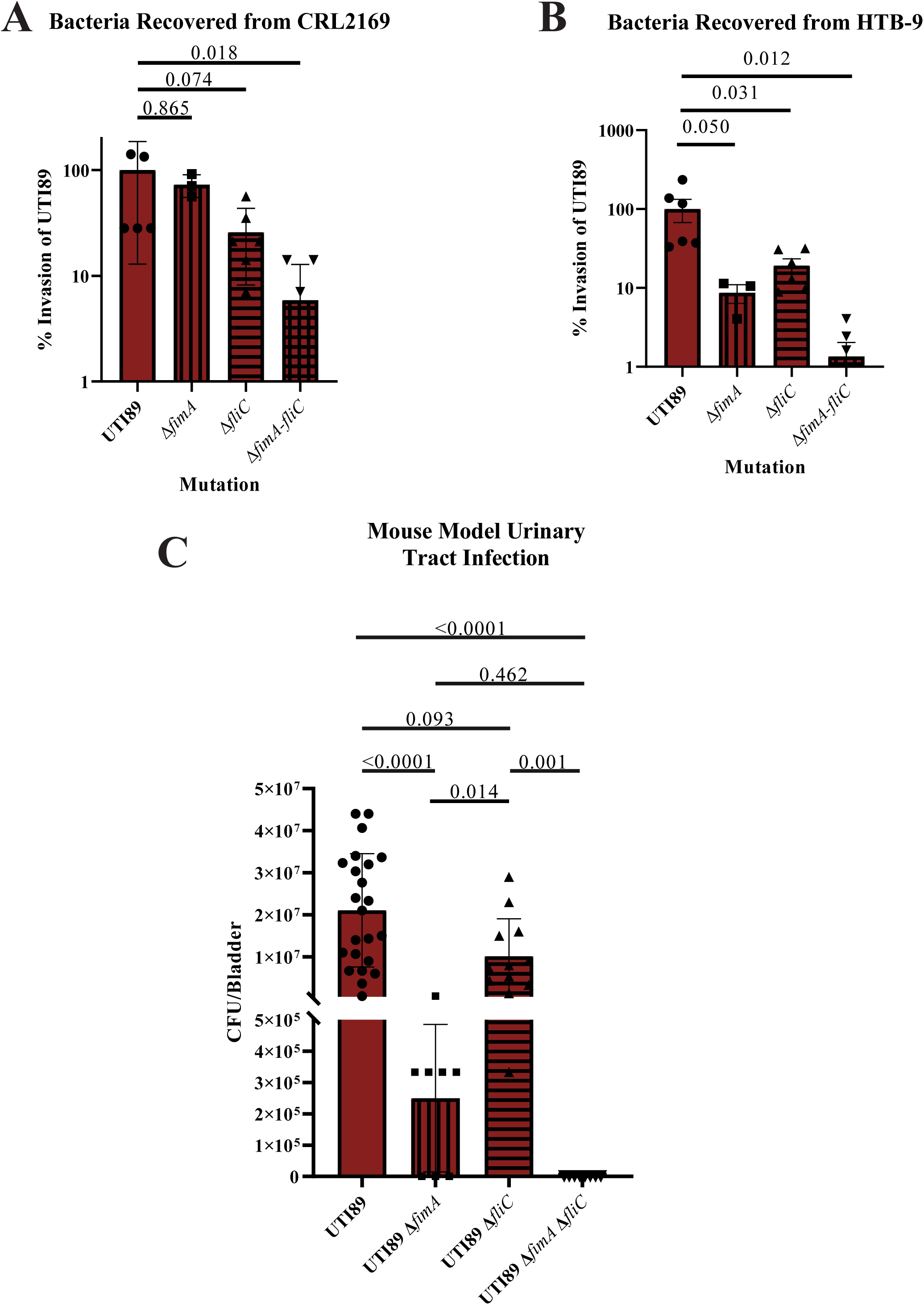
Comparison of wild-type UTI89 and mutants in the pili major structural subunit, *fimA*, the flagellar major subunit, *fliC*, and a double mutant in both *fimA* and *fliC*. Statistical significance was determined by Kruskal-Wallis non-parametric test of multiple comparisons. (A) Invasion into CRL2169. For parental, Δ*fimA*, Δ*fliC*, and Δ*fimA* Δ*fliC* strains, the samples sizes were 5, 3, 6, and 6, respectively. For the double mutant, three replicates had no recoverable bacteria. (B) Invasion into HTB-9. For parental UTI89 and Δ*fimA*, Δ*fliC*, and Δ*fimA* Δ*fliC* derivatives, the samples sizes were 6, 3, 6, and 6, respectively. For the double mutant, none of the six replicates had recoverable bacteria. (C) Mouse model of urinary tract infection. For parental, Δ*fimA*, Δ*fliC*, and Δ*fimA* Δ*fliC* strains, the samples sizes were 23, 8, 12, and 8, respectively. For the double mutant, none of the 8 replicates had recoverable bacteria.

Because many of the invasive strains express flagella, we tested whether a derivative of the non-invasive pili-dominant group A strain that expresses flagella instead of pili are invasive. We had previously isolated a W3110 derivative, called J1, that was hypermotile and expressed flagella instead of pili because of an *IS*5 insertion sequence in the *flhDC* promoter region (34). Although transcriptomic results suggest that J1 expresses pili and flagella equally (Fig. 2C), electron microscopy showed that J1 had flagella and no pili (34). Like its parent, J1 was non-invasive which indicates that flagella are not sufficient for invasion into CRL2169 (Fig 4).

**Figure 4.**
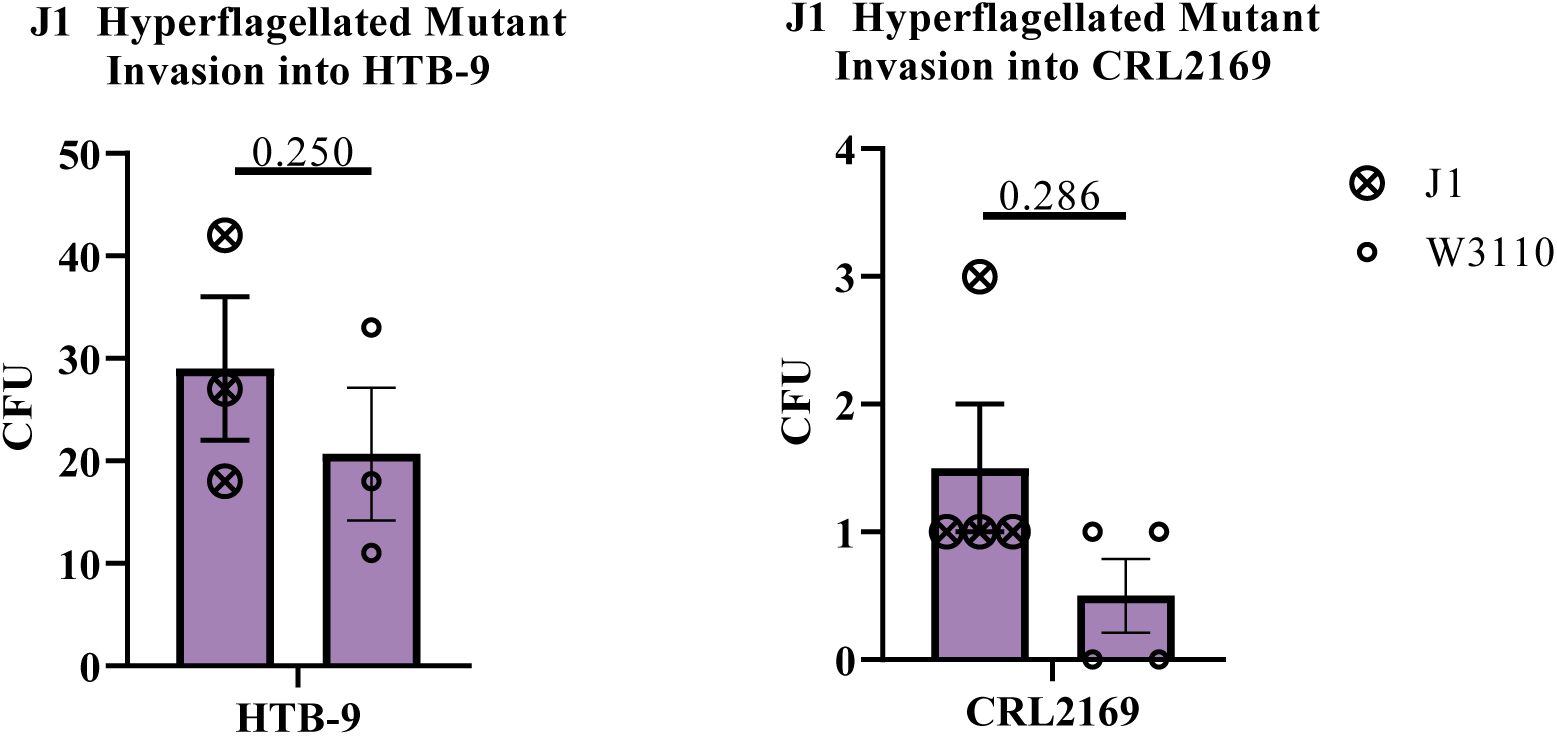
Invasion of a hyperflagellated derivative of W3110 in HTB-9 and CRL2169. No bacterial strain was considered invasive. ns, not significant

### CRL2169 is more differentiated than HTB-9 based on differences in endocytic receptors and the endocytic mechanism

Endocytic pathways initiated by flagella and pili form clathrin- and caveolin-dependent vesicles, respectively (Fig 5A). CRL2169 and HTB-9 are derived from well-differentiated grade 1 and moderately differentiated grade 2 carcinomas, respectively, which should correlate with several properties associated with binding and endocytosis. Greater urothelial differentiation is correlated with greater uroplakin expression (35), more TLR4 receptors (36), less active TLR2 receptors (37), and less endocytic activity (31, 32). Differences in expression of many of these genes between CRL2169 and HTB-9 can be determined from the Cancer Cell Line Encyclopedia (38, 39). CRL2169 had more transcripts for multiple pili-binding uroplakins (Fig 5B) which is consistent with a more differentiated state. CRL2169 had 23-fold more transcripts for TLR4, 1.7-fold more transcripts for TLR5, and 29-fold lower transcripts for TLR2 (Fig 5C) (38). We confirmed these differences in our cells by reverse transcriptase quantitative PCR (Fig A2). TLR4 levels are lower in bladder cancer cells, and TLR2 is associated with tumor progression (36, 37), which means that these results show that CRL2169 is more differentiated than HTB-9. HTB-9 has more transcripts for two dynamin alleles (DNM1 and DMN2), and summing the transcripts for both alleles shows that HTB-9 has twice the transcripts for dynamin (transcripts for DNM2 are more abundant) (Fig 5D). In sum, these results are consistent with differences in the differentiation state and endocytic pathways.

**Figure 5.**
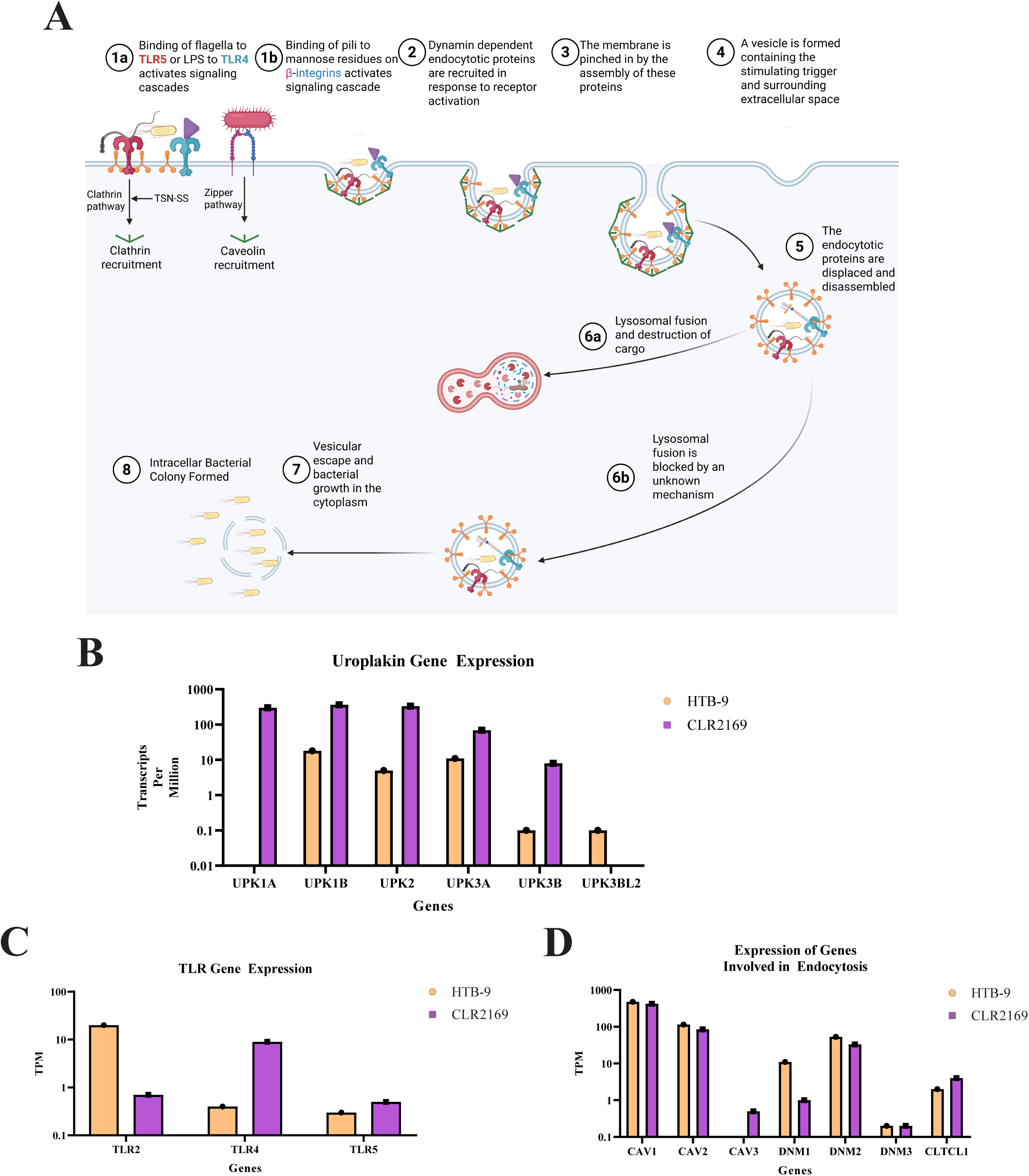
Overview of endocytosis and IBC formation and ratio of endocytic proteins and receptors. (A) Overview of endocytosis. Red and blue receptors are TLR5 and TLR4, respectively. Both TLRs are shown binding their activating signals, flagella and LPS, denoted by the black flagella and purple triangle. Green angles represent both clathrin and caveolin. Image was generated using Biorender.com. (B) Uroplakin transcript levels between CRL2169 and HTB-9 from public databases. (C) TLR-2, -4, and -5 transcript levels from public databases. (D) Transcript ratios for endocytosis genes between CRL2169 and HTB-9 from (38, 39).

Following this line of reasoning, we analyzed the effect of the endocytosis stimulator tanshinone II A (TSN) which stimulates the dynamin-requiring clathrin- and caveolin-dependent pathways for the hyperflagellated group B2 strain ECOR51 (40). ECOR51 was selected because it had the highest, most reproducible invasion frequency. TSN increased invasion into HTB-9 four-fold (p = 0.029) but had little effect on ECOR51 invasion into CRL2169 (Fig. 6). These results show a contribution of a dynamin-dependent pathway for flagella-dependent invasion into HTB-9, but not into CRL2169. The implied difference in endocytic pathways is consistent with greater differentiation of CRL2169.

**Figure 6.**
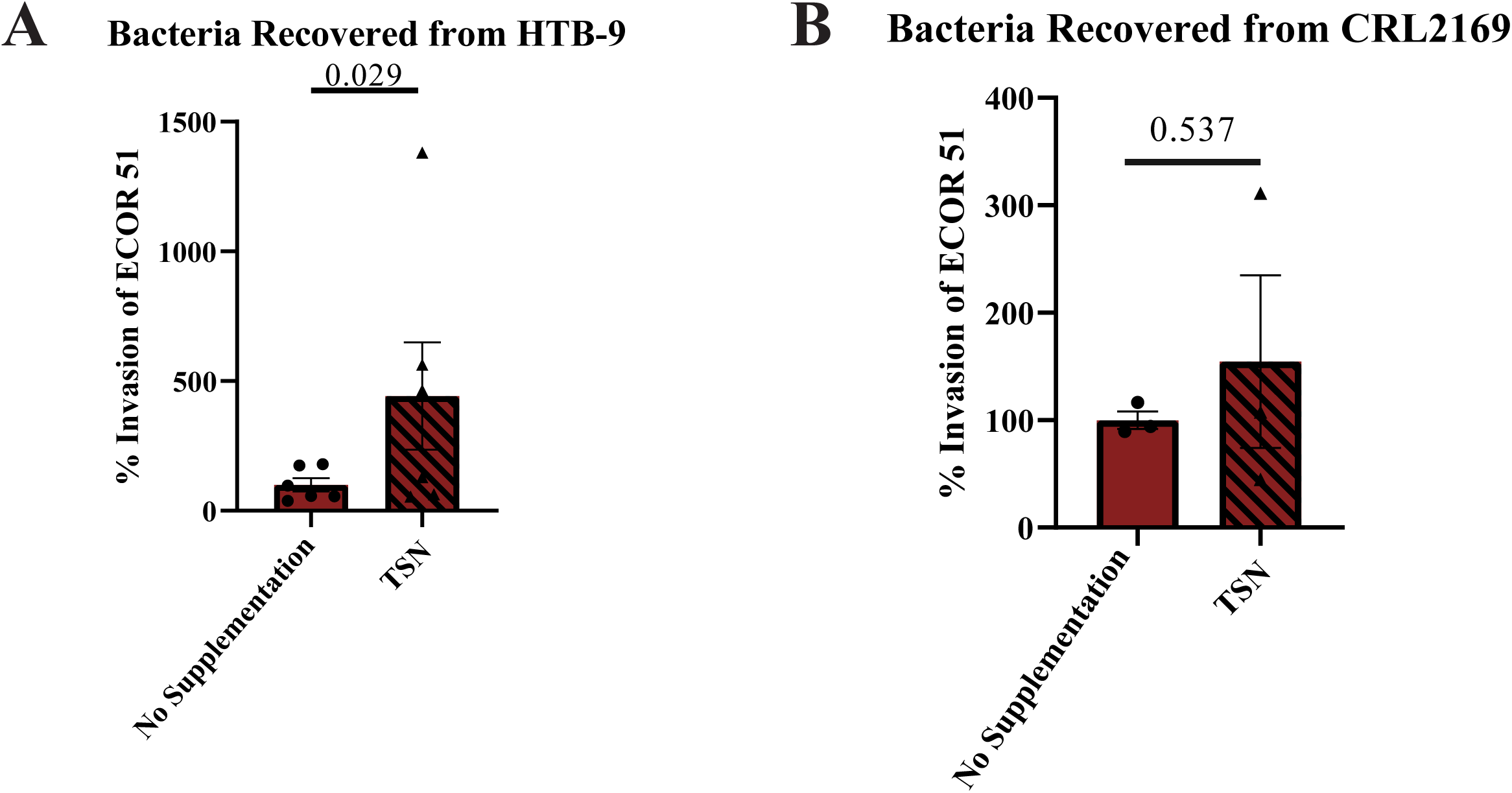
Chemical activation of endocytic pathways with tanshinone. Tanshinone (TSN) was added to human cell cultures two hours prior to the addition of the hyperflagellated strain ECOR51. Statistical significance was determined by a Kruskal-Wallis non-parametric test of multiple comparisons. ns, not significant.

## Discussion

We showed that phylogenetic group B2 strains invade CRL2169 cells better than strains of other phylogenetic groups. B2 strains are more prevalent during UTIs, but few relevant B2 properties are known. One important property is that B2 strains grow faster in urine than strains of other groups (41–43). Even though not nutrient rich, urine supports extremely rapid UPEC growth (41, 44). Consistent with rapid growth, UPEC strains have a rapid growth transcription program (45): B2 strains generally have more transcripts for genes associated with rapid growth, e.g., those for ribosomal proteins, when grown in a nutrient-rich medium (16).

Flagella were unexpectedly found to be important for invasion of B2 strains into CRL2169, even for the pili-dominant UTI89. The B2 strain ECOR53 did not invade CRL2169, which is the exception that proves the rule: ECOR53 has a low level of *fliC* transcripts. For the non-B2 strains, *fliC* transcript levels are low, which is expected because the medium glucose should suppress cyclic-AMP and flagella synthesis (46), but even a hyperflagellated strain in which this regulation is subverted did not invade. The anomaly is flagella expression in the B2 strains, which glucose should suppress, and which could result from either a B2-specific gene, altered regulation of cyclic-AMP synthesis, or both.

A major difference between the cell lines is that CRL2169 and HTB-9 cells are derived from grade 1 and 2 carcinomas, respectively: grade 1 carcinomas are more differentiated. Dedifferentiation increases endocytosis pathways (32), which is consistent with tanshinone-stimulated endocytosis into HTB-9, but not into CRL2169. A possible flagella-dependent tanshinone-independent entry mechanism into CRL2169 cells is a process that utilizes components of compensatory endocytosis which is a normal process of membrane internalization during bladder voiding (17, 47). The role of pili during UTIs is well established, although a subtle role for flagella has been acknowledged (17, 48). Our results with the relatively dedifferentiated HTB-9 are consistent with previous results that suggest the importance of pili for binding and intracellular invasion.

The novelty of our observations is the requirements for invasion into a relatively differentiated cell line, CRL2169. The importance of flagella may have been underestimated because of the conditions for cell culture assays and mouse infections. For binding assays with cultured cells, the glucose-containing medium should suppress flagella synthesis, and favor pili-dependent invasion. For mouse infections, the ∼2.0 M urea in the mouse bladder, which is >six-fold higher than in human urine (49), could relatively weaken or eliminate flagella binding and favor pili binding because of weaker binding of flagella to the epithelial layer compared to binding of pili. Flagella binding to TLR5, the presumed receptor, is at least two orders of magnitude weaker than pili binding to mannosylated proteins (deduced from results presented in (50, 51)). Our results support this possibility: for UTI89, flagella are more important for invasion into CRL2169, whereas pili are more important for a mouse infection. A further complication to unraveling the relative roles of pili to flagella is that physiological bladder conditions impair pili expression and function (52), although pili-dependent adhesion increases both pili expression and binding affinity (53, 54).

Limitations of past studies make it difficult to determine if the group B2 invasion skew and the role of flagella are relevant for UTIs. These limitations include overreliance on non-representative highly passaged “model” bacterial strains (16), a dedifferentiated, easily invaded cell line, HTB-9, and a mouse model system with an extremely high urea concentration that may require pili-dependent invasion. Invasion is a rare event, even in the absence of a glycosaminoglycan layer and host immunity, which is the situation for invasion studies with cell cultures. If differentiated urothelial cells are not invaded, then injured or partially defective cells may be invaded. Pathoadaptive isoforms of pili and flagella are known (15, 25), which suggests that some strains could rely primarily on flagella. If differentiated urothelial cells are invaded, then there is a phylogenetic skew for B2 strains, and the role of flagella has been underestimated.

## Materials and Methods

### Human cell lines and culture practices

The grade 2 male urothelial carcinoma cell line, HTB-9 (also called line 5637, ATCC-5637) and the grade 1 female urothelial carcinoma cell line, CRL-2169 (ATCC-SW 780) were grown in Lebovitz’s 15 (L-15) (ATCC-30-2008) medium supplemented with antibiotic-antimycotic (Sigma-A5955) and 10% FBS (Thermo-A3160401) at 37°C to roughly 80% confluency. Cells were split into 24 well plates, ∼50,000 cells per well, and grown again to 80% before infection. The endocytosis activator tanshinone, when added, was administered at 100 µM for 2 hours before introduction of bacteria.

### Bacterial cell lines and culture practices

Strain characteristics are described in Table 1. W3110 is a lab strain. The other strains are from a variety of sources as described in Table 1 (55–58). UTI89 derivatives were constructed as described (34). Strains were divided into the Clermont phylogenetic groups following the method described in (59).

To ensure that the bacteria were in a logarithmic phase of growth, bacterial cells were grown aerated for two hours in LB at 37°C. The authors note that overnight cultures or stationary cultures decreased invasiveness in comparison to logarithmic growing cultures (data not shown).

### Invasion assay

Cell lines were grown in 24-well plates to 80% confluence, media removed, and fresh L-15 supplemented with 10% FBS without antibiotics was added. Bacterial cell lines were grown to a logarithmic phase, then infected at a multiplicity of infection of 10:1. After a 2-hour incubation and medium removal, the infected cell lines were washed three times with Dulbecco’s PBS-A. Fresh L-15 media containing 100µg/mL gentamycin (biobasic-GB0217) was added, and the cells incubated overnight. The following day, the medium was removed, infected cell lines washed three times with Dulbecco’s PBS-A and suspended in 1/3 media volume of 0.1% triton-X 100 (biobasic-TB0198) in Delbecco’s PBS-A. The cells were placed on a rocking platform for 20 minutes at room temperature. Using a 1mL pipette tip, the cells were scraped off, and the triton-containing solution was plated on fresh LB plates. The plates were dried at room temperature then transferred to an incubator where the bacterial cells were grown overnight at 37°C. The next morning, Petri plates were imaged, and colonies enumerated. All data was done in technical and biological triplicate unless otherwise specified.

### RNA isolation and qRT-PCR analysis

Cell lines were grown to 80% confluence and lysed using a 24-gauge needle. RNA was isolated with a Qiagen RNeasy mini kit following the company’s protocol. DNA contamination was removed using DNase treatment, and the resulting RNA was subjected to a GeneJet RNA Cleanup and Concentration kit. To ensure the RNA was of high quality and quantity, Nanodrop and Qubit analysis was performed. RNA samples were ≥ 1 µg/µL with 260/230 ≥ 2.0, 260/280 ∼1.8, and RNA IQ scores of ≥ 8.

qRT-PCR employed TaqMan assay kits specifically for TLR-5, TLR-4, and ACTB. qPCR was performed on a Quantstudio 3 machine following recommended settings for the TaqMan assays. Relative quantity (RQ) scores were generated and normalized to β-actin (ACTB). Data was analyzed on the ThermoFisher cloud qPCR web application. Assays were done in technical and biological duplicate.

### Mouse infections

The mouse infections followed previously described procedures (33, 60). Bacteria were grown at 37°C in Luria-Bertani (LB) broth without shaking. Cultures were collected, their optical densities measured, and the bacteria resuspended in sterile PBS at 2×10^8^ colony forming units (CFU)/mL. For each bacterial strain, 10 female mice were employed and anesthetized by isoflurane inhalation. Once fully anesthetized (confirmed by hind paw pinches), urine was expelled by gently massaging and pushing on the bladder. A sterile urinary catheter was covered in lubrication and inserted into the urethra. A syringe containing 10^7^ CFU in 50 µL PBS was attached to the catheter, and the bacteria were slowly deposited into the bladder. The catheter was slowly withdrawn from the urethra, and the animal was returned to the home cage to recover. Mice were euthanized either 24 hours after infection, and the bladder and urethral tissue were harvested for further analyses. Transurethral PBS infusion was used as mock infection control. Bladders were harvested 24 hours post-infection, homogenized and serially diluted for CFU enumeration.

## Acknowledgements

The authors would like to thank Dr. Nikki Delk for her advice and guidance in cell culture techniques. The authors would also like to thank Dr. Tae Hoon Kim and the Kim lab for the use of their facilities. J.H. would like to thank Trusha Parekh, B.S., M.S. for her advice and guidance on cell culture techniques.

---

**Fig A1.**
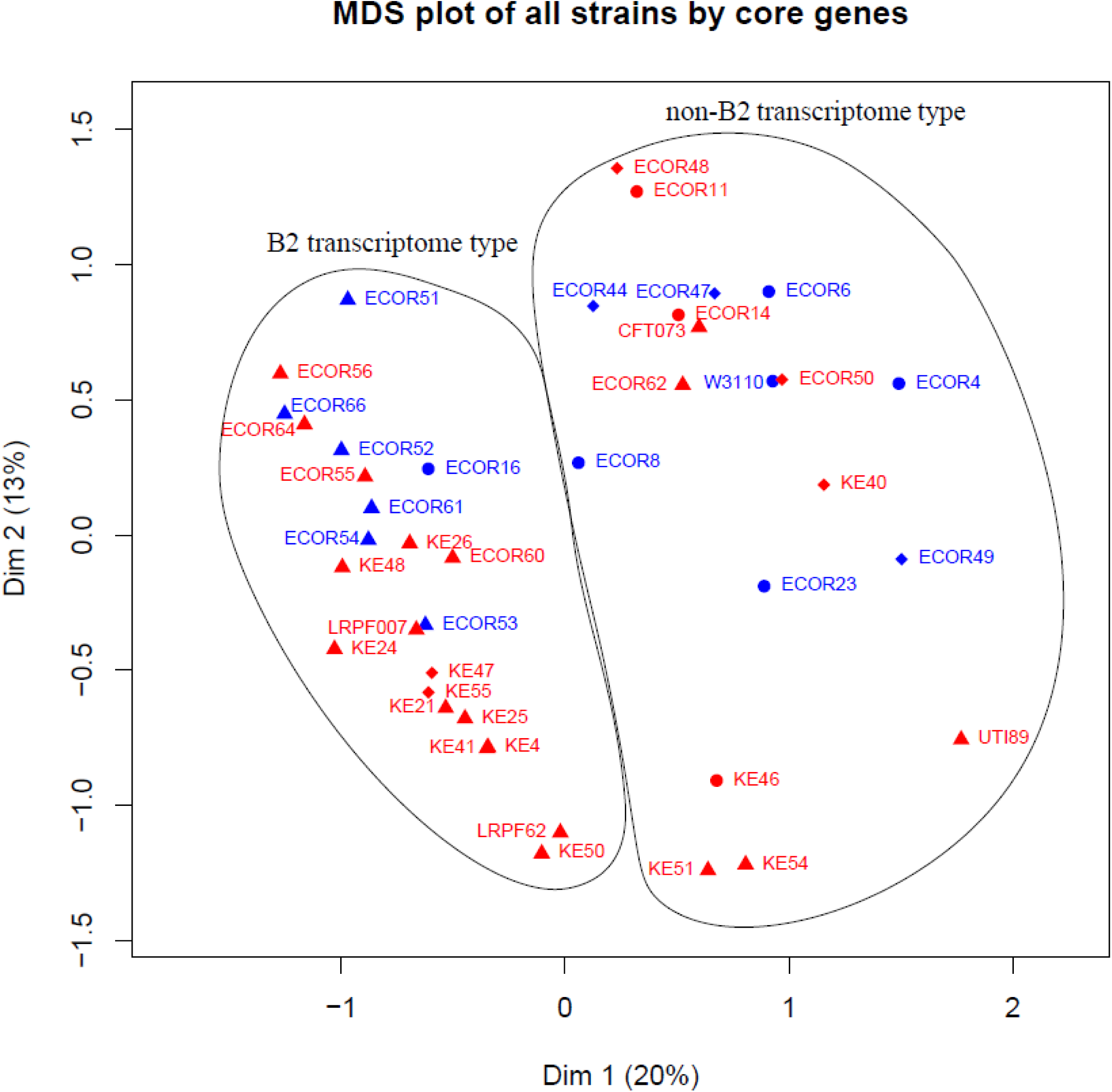
Transcriptome types updated from Hogins et al. (16). The transcriptome types were determined by K means clustering. The original plot had B2 and non-B2 types and four outliers that did not map within either cluster. In this update, all strains map into one of two clusters. Note that CFT073 and UTI89 map within the non-B2 cluster. The cells had been grown as described (16). Strains for phylogenetic groups A, B2, and D are indicated with circles, triangles, and diamonds, respectively. Uropathogenic strains, in red, were isolated from individuals with a urinary tract infection; non-pathogenic strains, in blue, were not isolated from urinary tract infection patients. No clear distinction exists between pathogenic and non-pathogenic strains.

**Fig A2.**
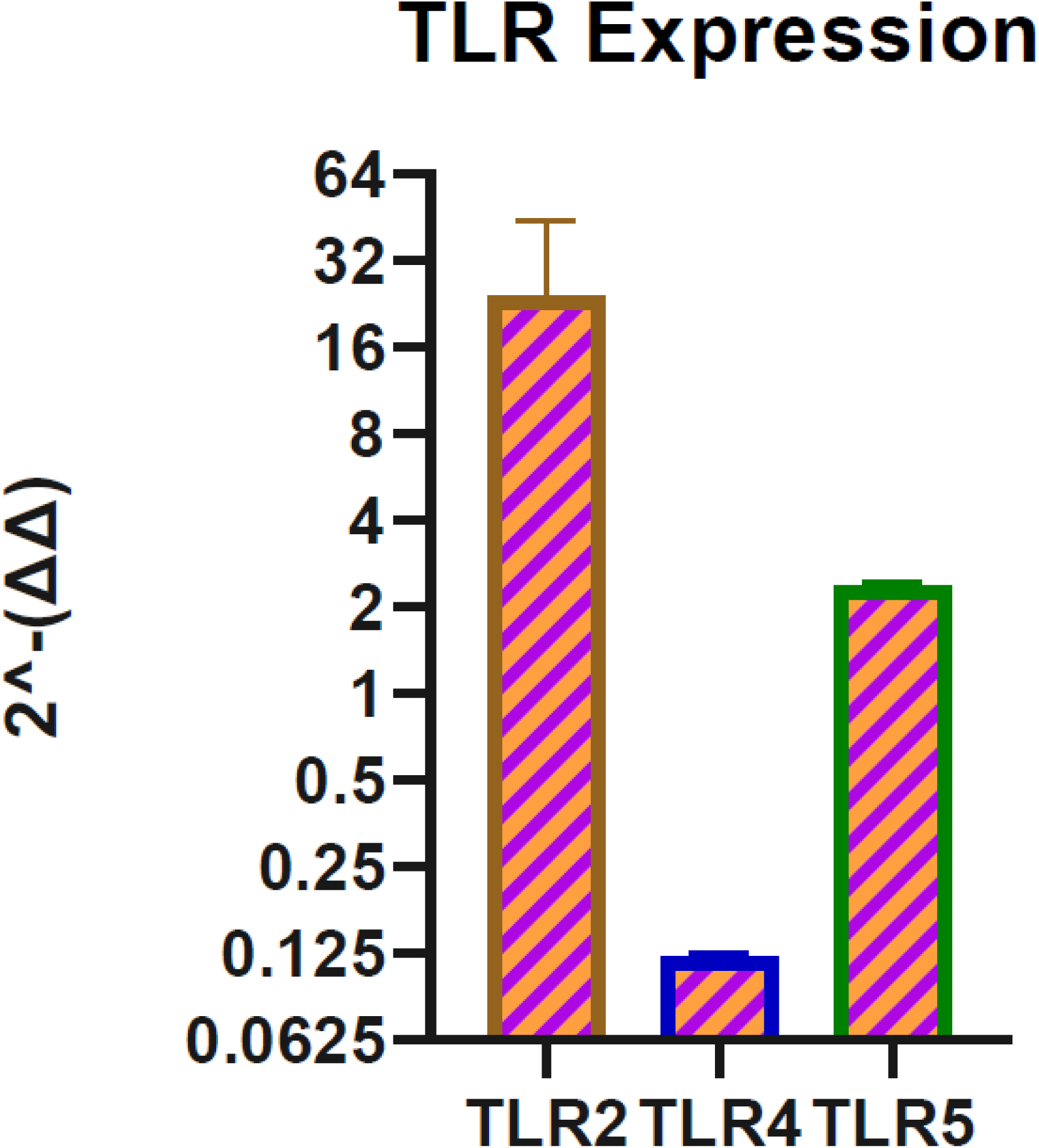
Transcript ratios of TLRs from HTB-9 to CRL2169 cells by reverse transcription-quantitative PCR. The results confirm that the TLR transcripts from CRL2169 and HTB-9 cells used in this paper correspond to previously published results.

